# Defecation by the ctenophore *Mnemiopsis leidyi* occurs with an ultradian rhythm through a single transient anal pore

**DOI:** 10.1101/511576

**Authors:** Sidney L. Tamm

## Abstract

Defecation in the ctenophore *Mnemiopsis leidyi* is a stereotyped sequence of effector responses that occur with a regular ultradian rhythm. Time intervals between repeated defecations of individual animals depend on body size, ranging from ~10 min in small larvae to ~1 hr in large adults. New features and corrections of previous reports of the gastrovascular system during and between defecations are described in detail by video microscopy. Contrary to the scientific literature, the defecating organ of the excretory complex is just one of the two anal canals which possesses the animal’s only anal pore. The anal pore is not visible as a permanent structure as depicted in textbooks, but appears at defecation and disappears afterward. DIC microscopy reveals that opening and closing of the anal pore resemble a reversible ring of tissue fusion between apposed endodermal and ectodermal layers at the aboral end. *Mnemiopsis* thus appears to have an intermittent anus and therefore an intermittent through-gut that reoccur at regular intervals. The temporality of a visible anal pore in *Mnemiopsis* is novel, and may shed light on the evolution of a permanent anus and through-gut in animals. In addition, mirror image dimorphism of the diagonal anal complex occurs in larval ctenophores but not in adults, indicating developmental flexibility in diagonal symmetry of the anal complex.

---

Many aspects of the functional morphology of ctenophores with regard to sponges, placozoans and cnidarians remain poorly understood, and are essential for understanding the early evolution of animals as integrated organisms (Dunn and Ryan, 2015). Evolution of the gut is a prime example. Contrary to a recent version of the historical record (Presnell et al., 2016), the prevailing consensus in both the scientific literature and textbooks is that the ctenophore gut is not a blind sac with a single opening (bottle-gut), but a through-gut with mouth and separate anus (Barnes, 1980; Brusca and Brusca, 1990; Bumann and Puls, 1997; Dunn et al., 2015; Franc, 1972; Hernandez-Nicaise, 1991; Hyman, 1940; Mayer, 1912; Main, 1928; Pearse, et al., 1987; Sullivan, 2010; Tamm, 2014a, 2016). This has been well known since the classic studies by Agassiz (1850) and Chun (1880), and was one of the reasons why Hyman (1940) set up Ctenophora as a separate phylum instead of remaining a subgroup of Coelenterata (now Cnidaria).

Nevertheless, many structural and functional features of the ctenophore gastrovascular system remain incompletely or incorrectly described. Here I report new features and correct past studies of the gastrovascular system in the ctenophore *Mnemiopsis leidyi*, and show that defecation occurs through only a single anal pore which appears and disappears with a regular ultradian rhythm.

## Background

Ctenophores are important members of the marine zooplankton (Haddock, 2007; Harbison, et al., 1978; Harbison and Madin, 1982). They have 180HHrotational symmetry in any plane through the oral-aboral axis (except for the anal complex), with the mouth at the front end and a sensory-motor statocyst or apical organ at the aboral pole (Dunn, et al., 2015; Hernandez-Nicaise, 1991; Tamm, 1982, 2014a, 2014b, 2015). Two orthogonal planes run along the oral-aboral axis: the sagittal plane in which the flattened elongated pharynx or stomodaeum lies, and the tentacular plane passing through the two tentacle pouches. Most organ systems are bilateral with respect to either the sagittal or tentacular plane.

Embryonic development and hatching leads to free-swimming larvae which, except in beroids, resemble the cydippid *Pleurobrachia*. When cydippid larvae of *Mnemiopsis* reach a size of ~5 mm long, two developing lobes appear in the sagittal plane on either side of the mouth, marking the transition from cydippid to lobate body plans. Progression to the lobate form continues with larval growth and typically is attained in 1.2 - 1.5 cm larvae (Tamm, 2012b). Later stages of growth are referred to here as young animals and adults.

The gastrovascular system of ctenophores consists of six distinct parts (see also Hyman, 1940; Main, 1928; Mayer, 1912): a muscular ciliated mouth with lips, a long flattened stomodaeum (pharynx) with muscular folds, a turtleneck-shaped esophagus with medial lappets, a centrally located stomach (funnel or infundibulum), a branching food canal distribution system, and an aboral canal-anal canal excretory complex. Each portion of the hexapartite gastrovascular system is anatomically and physiologically specialized to perform a distinct function in food processing.

Following ingestion of prey by various means in different orders (Colin, et al., 2010; Haddock, 2007; Harbison and Madin, 1982; Harbison, et al., 1978; Moss, 1991; Swanberg, 1974; Tamm, 1982, 2014a; Tamm and Moss, 1985; Tamm and Tamm, 1987, 1991; Waggett and Costello, 1999), consecutive phases of extracellular digestion take place in an oral-aboral sequence along the stomodaeum (Bulman and Pils, 1997). Bulky indigestible food fragments (exoskeletons, bones, etc) never enter the canal system and are transported backwards along the edges of the stomodaeum and ejected through the mouth (Main, 1928; Bulman and Pils, 1997).

The esophagus at the aboral end of the stomodaeum acts as a gastric mill or food processor to break up pre-digested prey. Its lateral shoulders are lined with hammer-headed compound cilia arranged in circular arrays on each side (Tamm, 2014a). Presence of food in the aboral region of the stomodaeum is usually accompanied by widespread activation and vigorous beating of the esophageal cilia, causing countercurrent whirling on opposite shoulders and maceration of the pre-digested prey into small food particles which are swept into the stomach by two medial ciliated lappets, which serve as filters as well as valves for food particle flow into the stomach.

Distribution of food from the stomach to various parts of the body, and detailed description of defecation of waste products by the anal complex, are reported in **Results**.

## Materials and methods

### Organisms

*Mneiopsis leidyi* A. Agassiz, 1856, were carefully dipped from the surface of the sea at Woods Hole, Massachusetts. Animals were used immediately for observations or kept briefly in seawater for observations later the same day. Free-swimming larvae and young animals in the plankton were collected from flat-bottomed glass bowls of fresh seawater under a stereomicroscope. Animals were not fed and were observed under standard laboratory conditions.

### Video microscopy

Individual ctenophores were pinned in the sagittal plane in a Sylgard-floored glass fingerbowl or petri dish of seawater and observed under a Wild stereomicroscope. The gastrovascular system was imaged with a phototube-mounted digital video camera connected to a VCR with a timer and frame/field counter allowing still field (17 ms) playback and printing of VHS tapes (Tamm, 2012a, b). Living transverse thick sections of adults containing the anal complex were dissected and placed aboral pole upward in a petri dish of seawater for stereomicroscopy. Larvae were observed in different orientations on Vaseline-ridged well slides of seawater and imaged by Zeiss DIC (Differential Interference Contrast) video microscopy.

### Analysis

Timing of gastrovascular events was determined by playback analysis of video tapes with an audio track and illustrated notes transcribed during observations. Intervals between repeated defecations of individual animals were timed to the nearest minute. The stereotyped sequence of gastrovascular effector responses preparatory to defecation provided an alert to begin video recording of the defecation process. The number of successive defecations followed in different animals varied, depending on the length of intervals and other factors.

## Results

### Interludes between defecations

Nutrient distribution to various parts of the body occurs during interludes between bouts of defecation. A previously undescribed feature of the esophageal cilia is their switching between activation or quiescence on opposite shoulders during interludes (Fig. 1). The function and physiological basis of this local control of esophageal ciliary activity is not yet known.

**Figure 1.**
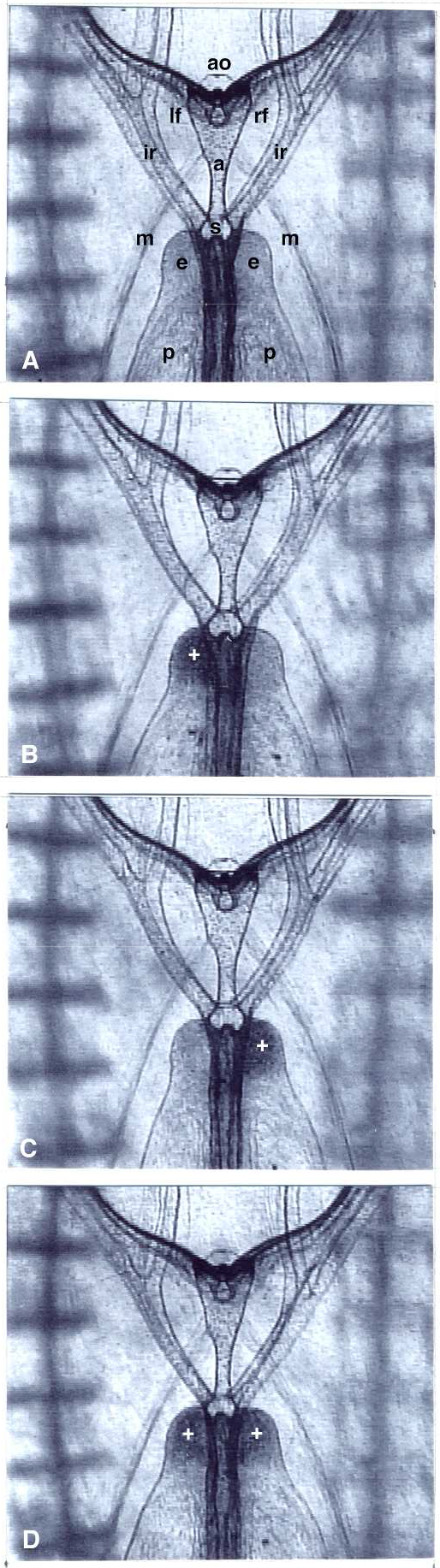
Patterns of activity of esophageal cilia during interlude in a medial sagittal view of a 2 cm young animal (aboral end up). Cilia on opposite sides of the esophagus (e) switch between quiescence (light gray) and active beating (white plus on dark gray), often independently (B, C). The dark structure running down the middle of the esophagus and pharynx represents the superimposed pair of paragastric canals and also pharyngeal folds. Ladder-like subtentacular comb rows are out of focus. Photographs are not arranged chronologically; time elapsed from A to D is 10 min. Note apical organ (ao), left and right anal forks (lf, rf), aboral canal (a), stomach (s), interradial canals (ir), pharynx (p), and retractor muscle bundles (m).

Food particles from the stomach exit into four interradial canals which run diagonally into the four quadrants of the body (Figs. 1, 2) and bifurcate into paired adradial canals. The adradial canals become the meridional canals running under the eight comb rows. Two paragastric canals run orally on either side of the stomodaeum (Fig. 1), and two tentacular canals supply the tentacle bulbs. The diameters of the endodermal canals and the directions of fluid and food particle flow driven by short cilia lining the canals are regulated and display various patterns of nutrient transport during interludes. For example, one-way transport of food particles in the interradial canals may be directed away from the stomach toward the periphery or in the opposite direction back into the stomach in various patterns that are coordinated (Fig 2). Transport of food particles in the meridional canals underlying the comb rows was observed to be bidirectional during interludes, confirming Gemmill (1918). During interludes most food particles in the stomach do not enter the aboral canal, or do so only briefly and return to the stomach. As a result, the aboral canal and anal canals remain almost empty of food particles between defecations.

**Figure 2.**
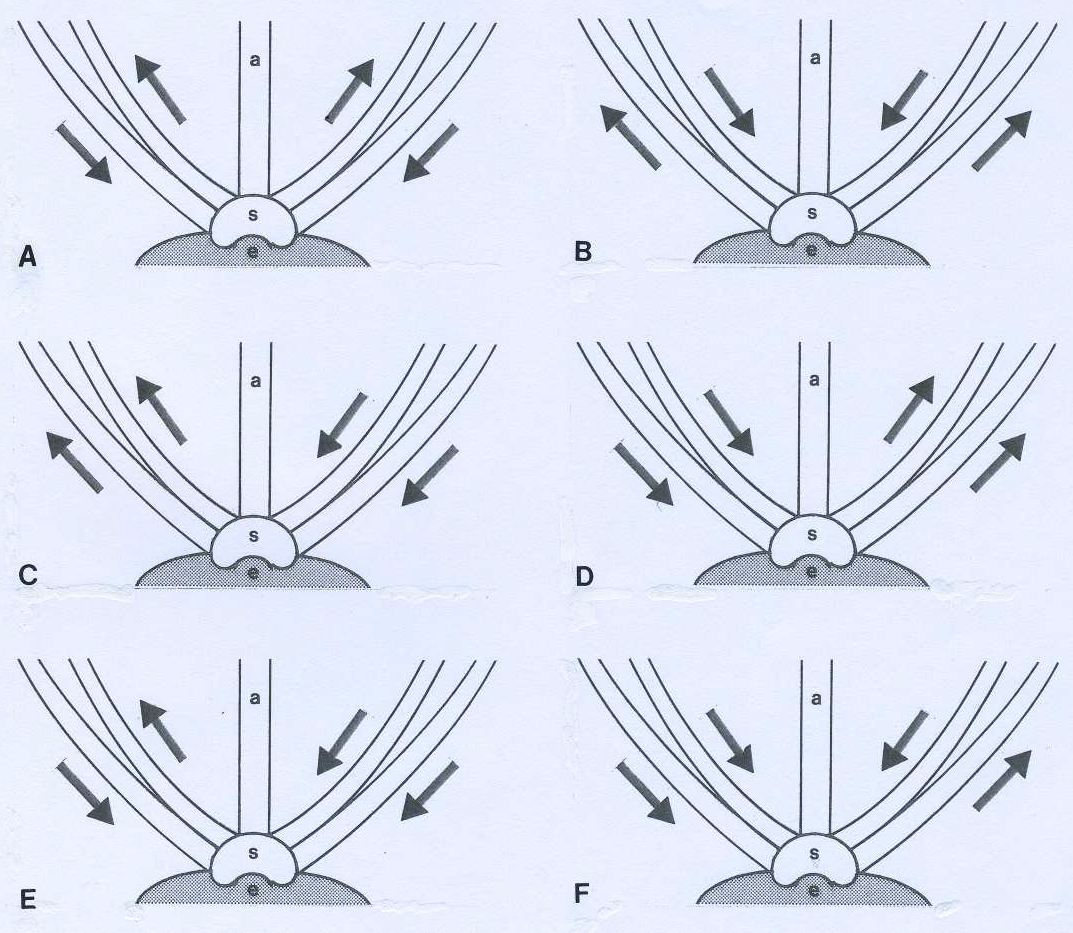
Diagram of coordinated patterns of one-way transport of food particles in the two pairs of interradial canals (arrows) during interlude, based on video analysis of a 1.2 cm young animal viewed in the sagittal plane. The panels are arranged by related patterns. The aboral canal (a), stomach (s), esophagus with medial lappets (e), and interradial canals are open to one another in spite of solid lines demarking junctions. A, B. Opposite directions of flow in each pair of interradial canals (into vs. out of the stomach). C, D. Direction of flow is the same in each interradial pair but opposite to the other pair. Particles go into or out of the stomach in opposing pairs. E, F. Three interradial canals have inward particle flow and one canal has outward flow. The outward interradial canal is different in E vs. F.

### Anal complex

In sagittal views of animals, the aboral canal extends along the oral-aboral axis from the stomach to the underside of the apical organ where it branches or forks into two opposing anal canals along the sagittal plane. Each anal fork is asymmetrically bi-lobed on either side of the apical organ. In sagittal views, the larger and smaller lobes of an anal canal are seen superimposed in each fork. The shorter lobe is blind and terminates in a dense cap (Fig. 3). One of the two opposed major lobes (in the right or left fork, depending randomly on which sagittal side is pinned against the Sylgard) appears larger and closer to the aboral surface than the opposite one during interludes (Fig. 3). It is this anal canal (or fork) which will later defecate.

**Figure 3.**
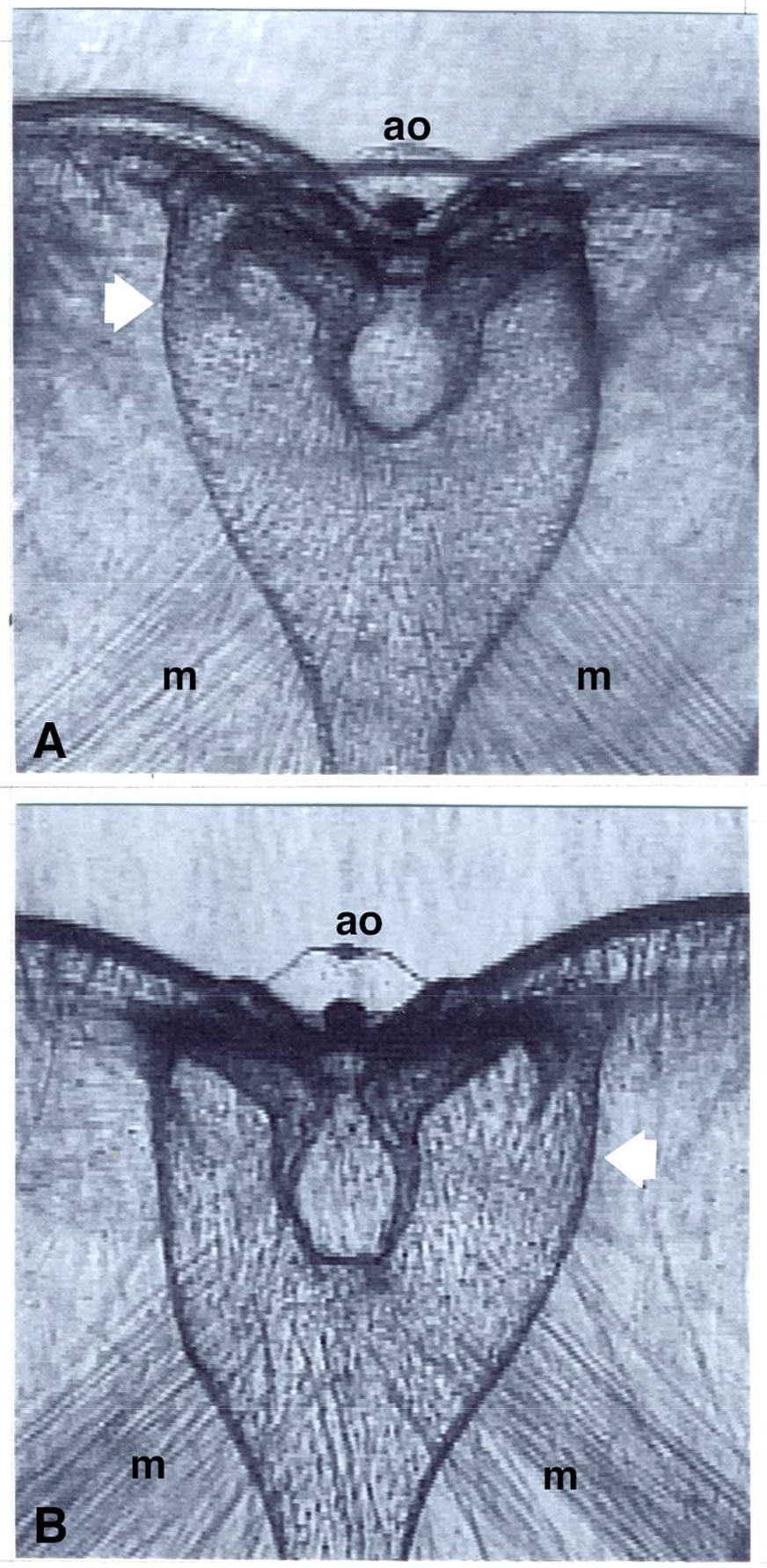
Interlude sagittal views of bi-lobed anal forks/canals on either side of the apical organ (ao) (aboral end up). The larger and smaller lobes of an anal canal are seen superimposed in each fork. The smaller blind lobes have a curved dense cap and are similar in size. Whether a given fork is left or right depends on random orientation (see text). m, retractor muscle bundle. A. 5 cm animal in which the major lobe of the left fork (white arrow) is larger and closer to the aboral surface than the major lobe of the right fork which cannot be distinguished. B. 4.2 cm animal in which the major lobe of the right fork (white arrow) is larger than that of the left fork.

Aboral views of the anal canals in larval and adult *Mnemiopsis* show the two major anal lobes opposite one another on a line diagonal to the sagittal and tentacular planes, while the two smaller capped lobes are opposed on the other diagonal. Again, the two major lobes are not the same size, and the larger one is used for defecation.

### Defecation

Periodic bouts of defecation interrupt and shut down food handling and distribution in the canal system. Video stereomicroscopic analysis of ctenophores of various sizes viewed in the sagittal plane (3 mm - 5 cm body length; n = 25) reveals that the earliest indication of an impending defecation is a change in shape of the stomach. The wide windshield-shaped stomach characteristic of interludes gradually narrows laterally into a rectangular box along the oral-aboral axis until it encloses the projecting medial lappets of the esophagus (Fig. 4). Before or during stomach narrowing, the field of esophageal cilia becomes predominantly or completely inactive. The onset of stomach narrowing was used as a visible signal to begin video recording of defecation, and is designated as time zero in the following account of sequential stages of defecation (Fig. 4).

**Figure 4.**
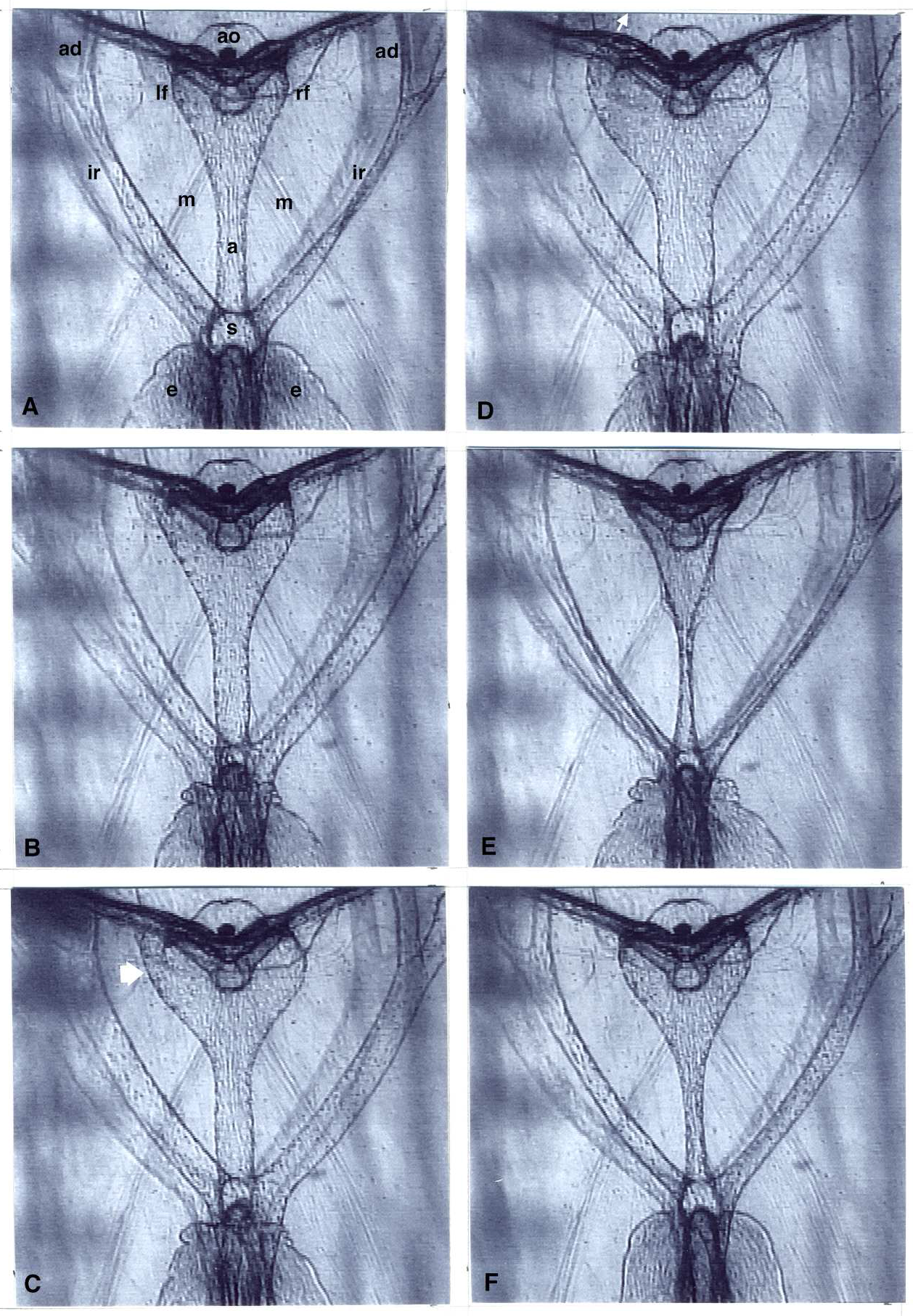
Main stages of defecation in a 1.2 cm young animal viewed in the medial sagittal plane (aboral end up). Note apical organ (ao), left and right anal forks (lf, rf), aboral canal (a), interradial canals (ir), adradial canals (ad), stomach (s), esophagus (e), retractor muscle bundles (m) and out-of-focus regions of subtentacular comb rows. A. Configuration at the end of an interlude, just before the onset of stomach narrowing. The left anal fork (lf) is wider than the right fork (rf). Some cilia in the esophagus are actively beating (darker central regions). B. 2 min later: all canals are swelling and the stomach has narrowed laterally to enclose the medial lappets of the esophagus. The contracting shoulders of the esophagus are crumpling and the cilia are quiescent (light gray). C. 3 min later: the canals have swollen further, and the major anal lobe of the left fork (white arrow) is expanded more than the major lobe in the right fork and is close to the aboral surface. The esophagus is constricted, blocking further entry of food into the narrowed stomach. D. 3.5 min later: the major anal lobes and all canals have reached their maximum volume. The greatly swollen major lobe of the left fork has pushed up the aboral surface and is now defecating (white arrow). E. 5.5 min later: both anal forks and all canals are greatly narrowed, and defecation has ended. Esophageal walls remain corrugated and their cilia are inactive. F. 8.5 min later: the anal forks, all canals and the stomach have widened, approaching the interlude configuration. The esophageal walls have relaxed and are smooth, but the cilia are not yet beating. The retractor muscle bundles do not shorten or change their appearance in the figure during swelling or constriction of the anal forks and canals.

At 1-2 min, the aboral canal and anal canals, as well as the interadials and adradials, begin to enlarge in diameter (Fig. 4 B - D). This widening of canals is not due to contraction of the two fan-shaped bundles of muscles extending orally from under the apical organ, as reported by Presnell, et al. (2016). To the contrary, these prominent groups of smooth muscles (Figs. 1, 3, 4) arise from a hub beneath the thickened epithelial floor of the apical organ and terminate orally in the mesoglea, not on canal walls, where they function to retract the apical organ into the body during avoidance responses to mechanical stimuli.

2-3 min after the onset of stomach narrowing, the smooth shoulders of the esophagus begin to constrict and become increasingly crumpled or corrugated (Fig. 4B; see Main 1928), effectively blocking further entry of food particles into the narrowing stomach. DIC microscopy of small animals in slide preparations shows that widening (and later narrowing) of the endodermal canals, as well as narrowing (and later widening) of the stomach, and crumpling (and later relaxation) of the esophageal walls, are not associated with shortening or extension of any mesogleal muscles.

At ~ 3 min, the bidirectional flow of particles in meridional canals characteristic of interludes becomes directed centrally into the stomach via adradial and interradial canals, where circulating particles now begin to enter the aboral and anal canals. The meridional canals constrict greatly in diameter and are soon emptied of particles. The interradial canals, and particularly the aboral canal and major lobes of the two anal canals, continue to swell (Fig. 4 B, C), becoming densely filled with circulating waste particles. The larger major lobe of one anal canal (in the left fork or right fork of a given animal, depending on random pinning of sagittal sides), expands more than the major lobe on the other side, and moves gradually closer to the aboral surface, which it pushes up as a rounded protrusion (Fig. 4 C, D; Fig. 5). The endodermal tip of this major lobe becomes closely apposed to the overlying ectodermal surface, excluding the intervening mesoglea. At the maximum volume attained by the greatly swollen aboral canal and protruding major anal lobe (at 3-4 min, Fig. 4 D), defecation suddenly begins as a stream of waste particles exiting the newly appearing anal pore at the tip of the protruding anal lobe (Fig. 4 D; Fig. 5).

**Figure 5.**
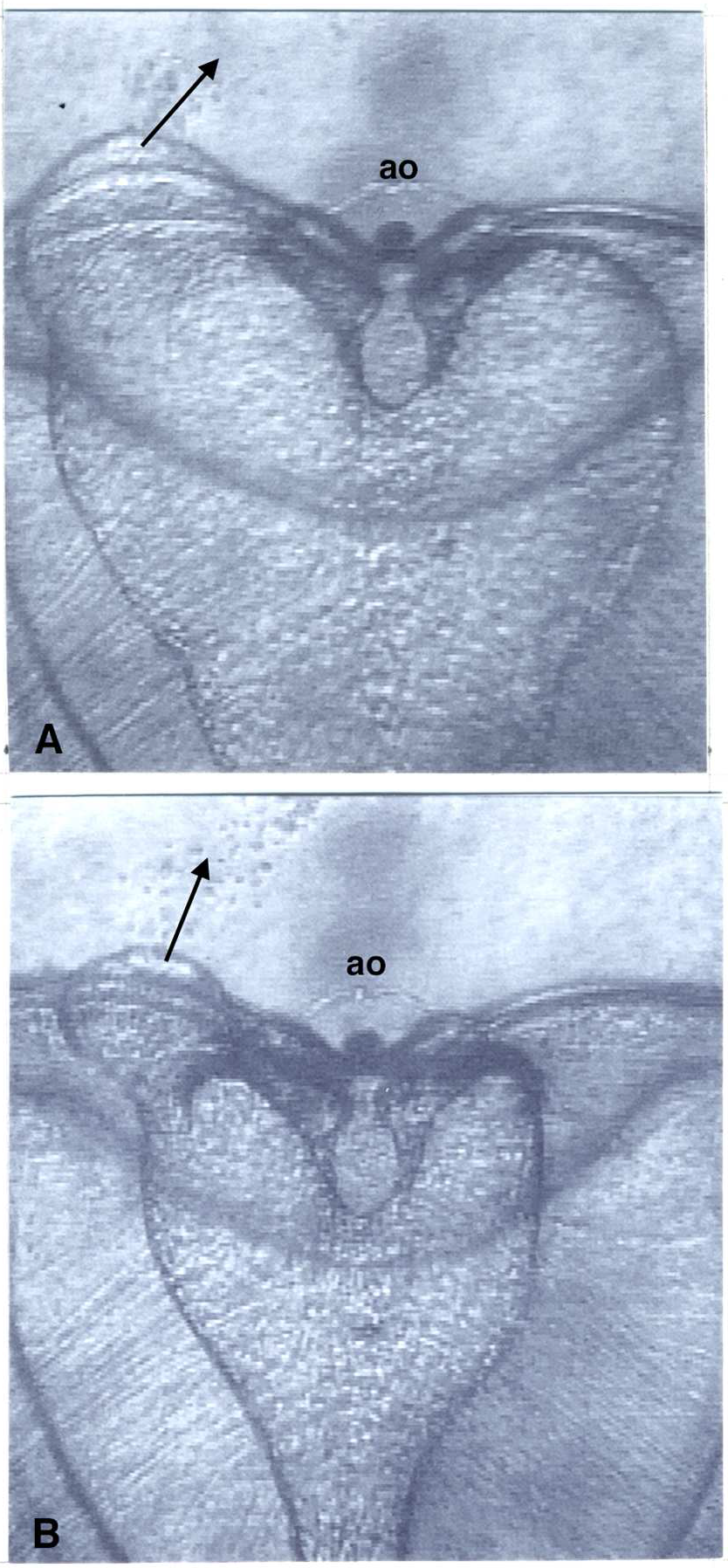
Successive stages of defecation by the greatly swollen major anal lobe in the left fork of a 5 cm adult animal viewed in the sagittal plane (aboral end up). Retractor muscles (diverging fibers) originate under the apical organ (ao). A. Maximum volume of the major anal lobes in the forks. The larger lobe in the left fork pushes up the aboral surface. Waste particles are expelled from the visible anal pore (arrow). B. 3.8 min later: both major anal lobes and the aboral canal have narrowed, revealing the densely capped blind lobe in each fork. The left major lobe is still protruding and its open pore continues to defecate (arrow).

Approximately 5-10 s after defecation starts, the aboral, anal and interradial canals begin to narrow in diameter. Circulating particles remaining in the non-defecating major anal lobe are swept into the defecating one and expelled through its pore. At this time one or both anal canals/forks sometimes undergoes rapid low-amplitude fluctuations in diameter for several minutes during waste expulsion. These anal canal pulsations always occur in the defecating anal canal/fork, and usually in the non-defecating one as well. In cases where both anal canals/forks pulsate, the pulsations in the defecating one are usually stronger and more frequent than in the non-defecating one. These vibratory movements may facilitate discharge from the defecating anal canal and fork. The canals and anal forks become greatly constricted at the end of defecation (Fig, 4E) and then slowly widen to the interlude configuration (Fig. 4F).

Since waste products entering the stomach and aboral canal come from all four interradials on both sides of the sagittal plane, a waste particle or clump travelling in a given interradial may be excreted from the anal canal/fork on that side (ipsilateral) or the one on the opposite side (contralateral), depending on the side of the defecatory anal canal/fork. Clumps of incoming waste particles reaching the base of an interradial always make a sharp turn entering the narrowed stomach, and commonly proceed into the aboral canal initially on the same side as they entered. If the defecating anal canal/fork happens to also lie on this side (ipsilateral), the waste clump continues straight up the aboral canal into the anal canal/fork and is expelled. In contralateral cases, however, waste clumps veer across the aboral canal and are swept into the defecating anal canal/fork on the opposite side (Fig. 6). This novel and sophisticated example of waste trafficking is presumed to be driven by a localized change in ciliary activity that diverts the waste stream into the defecating anal canal.

**Figure 6.**
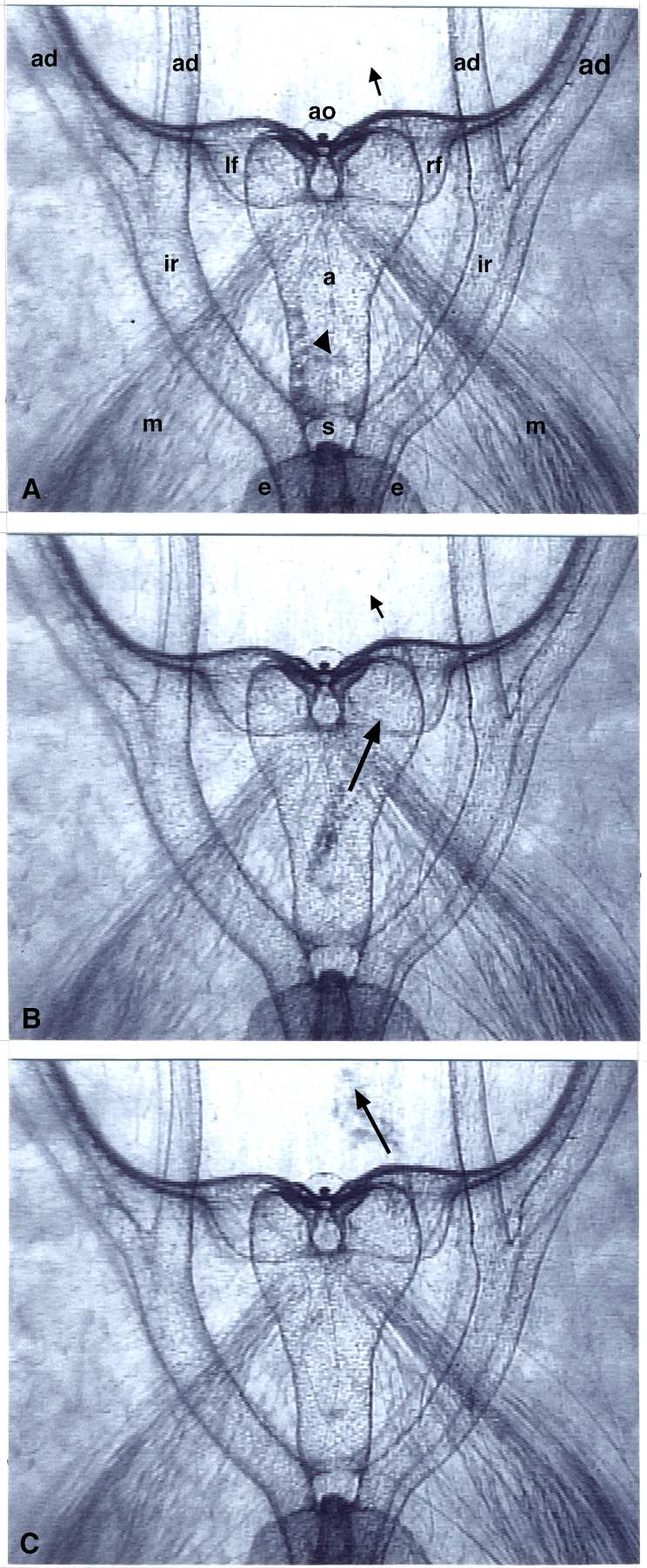
Contralateral excretion of waste clumps in a 4.2 cm adult animal viewed in the sagittal plane (aboral end up). Note apical organ (ao), left and right anal forks (lf, rf), aboral canal (a), stomach (s), interradial and adradial canals (ir, ad), esophagus (e), and retractor muscle bundles (m). The major anal lobe of the right fork is greatly swollen and has pushed up the aboral surface during defecation (arrows). The capped blind lobe in each fork is also visible under the aboral end. A. A train of waste clumps from a left interradial canal had turned through the stomach and is stalled briefly along the left side of the aboral canal (black arrowhead). B. The waste train slants across the aboral canal towards the enlarged defecating right fork (arrow). C. The waste train is expelled through the right fork (arrow).

Waste products, particularly large clumps or train-like aggregates, do not always pass uninterruptedly through gastrovascular canals, but sometimes stop briefly and then resume their journey out through the anal canal. Several instances of temporarily stalled waste trains have been observed in interradial canals and aboral canals (Fig. 6). The fecal masses may rotate or circle in place, or simply remain stationary. Local changes in ciliary activity underlying such stop-and-go waste removal have not yet been imaged. Near the end of a defecation, as the aboral canal and anal canals become progressively more constricted, any large fecal bolus exiting from the stomach is accompanied by a propagated widening of the aboral canal wall around the bolus, which is subsequently excreted. Such temporary increases in canal diameter by large propelled cargo illustrate the flexibility and elasticity of the endodermal wall to elevations of internal pressure.

### Anal pore appearance and disappearance

Aboral views of four 2 - 4 mm long larvae by DIC video microscopy show that repeated defecations by a larva always involved the same major anal lobe. The aboral surface of a swollen, about-to-defecate major anal lobe appears uniformly smooth with no visible structural indication of its future pore (Fig. 7A). As more waste products accumulate and circulate in the enlarged lobe pressing against the abutting ectodermal tissue, a tiny circular opening suddenly appears on the outer surface. As the opening increases in diameter, underlying waste particles exit from the lobe in a steady stream. Larger clumps of fecal matter remain in the lobe until they fit through the expanding pore and are expelled (Fig. 7B, C). No structural changes in the rim of the pore or the surrounding surface are evident during opening of a pore, nor are any indications visible of a ring-like sphincter imagined by Presnell et al (2016).

**Figure 7.**
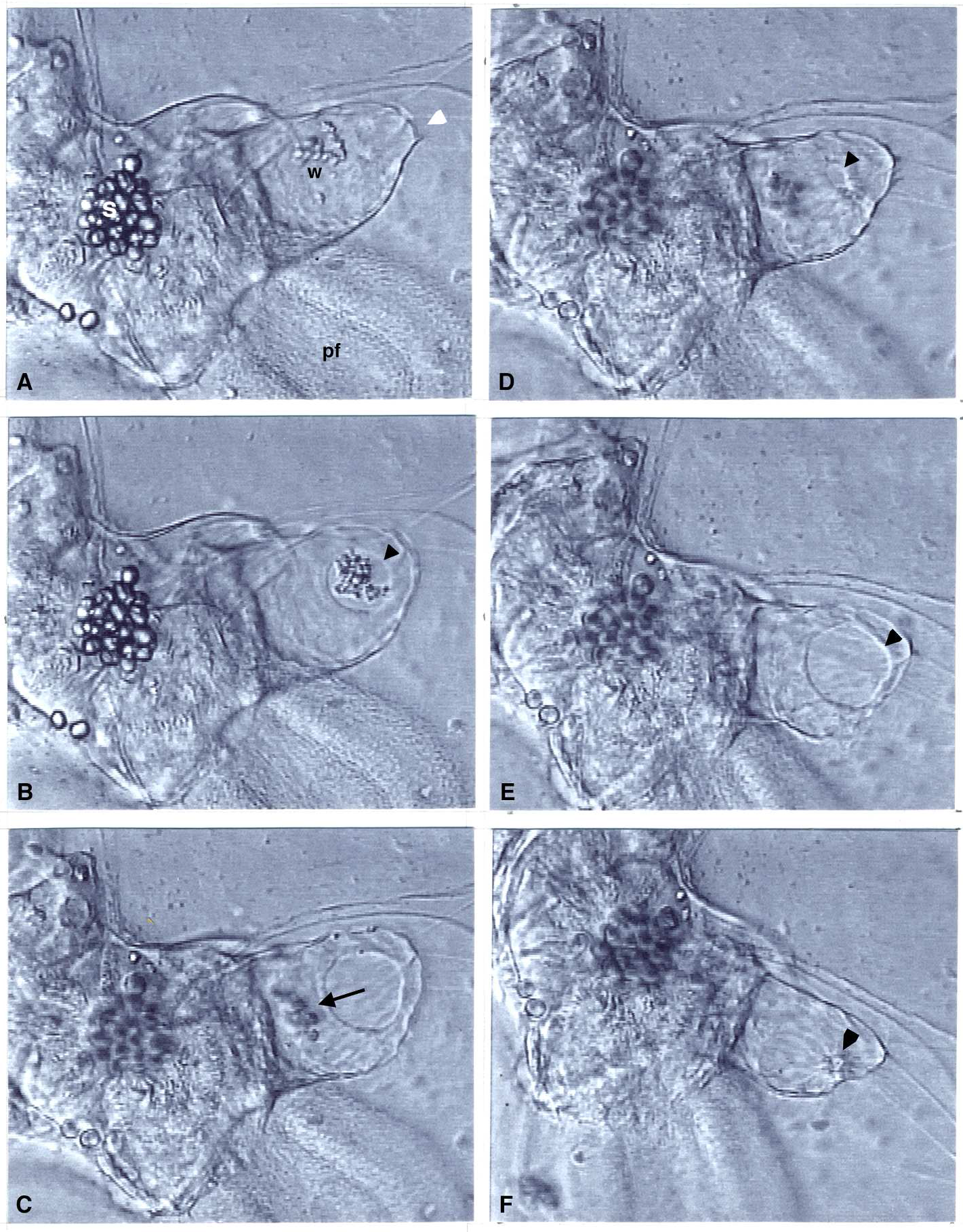
Anal pore opening and closing during the first 2 cycles of a 4 cycle defecation by a 0.18 cm larva (aboral DIC view). The cellular statocyst (s) lies at the center of the bow-tie shaped opposing bi-lobed anal canals. The swollen defecating major anal lobe (white arrowhead) is at the top right of the positive diagonal (see Fig. 8A). A polar field (pf) is visible on one side of the sagittal axis. A. Accumulated waste particles (w) circulate in the swollen major anal lobe before the anal pore appears. B. The pore opens (arrowhead) over the waste clump. C. The waste particles (out of focus) are expelled (arrow) from the enlarging pore. D. The pore closes partially (arrowhead). E. The pore enlarges again (cycle 2) to a maximum diameter (arrowhead). F. The pore closes completely (arrowhead). Note the central dense body surrounded by radiating lines (arrowhead). This closed structure later disappears. The larva slowly rotates under the coverslip in E and F.

Six of the 16 defecations (~38%) observed in the four larvae involved a single opening and closing of a pore (one cycle). Four defecations (25%) comprised two successive cycles of opening and closing of a pore, three defecations (~19%) involved three cycles of opening and closing of a pore, and three defecations (~19%) involved four cycles.

A pore may open to variable diameters in successive defecations or between cycles of a single defecation, and fluctuate in size within a cycle. A pore may also pause and remain open for variable times. Opening or closing of a pore may proceed smoothly or with intermittent pauses. In addition, the initially circular outline of a pore may be perturbed into oval or more irregular shapes by changes in lobe configuration during defecation.

Maximum diameters of fully opened pores in the 2 - 4 mm larvae were usually ~ 40 μm. During successive defecations (up to seven) of a single larva, the maximum diameter of fully opened pores varied. During defecations with multiple cycles, the maximum open pore diameter varied between cycles, and were usually smallest in the last cycle.

Closing of a pore begins as a decrease in its diameter without apparent structural changes at its edge or on the surrounding surface. As the pore becomes very small, it becomes surrounded by lines or folds of the surface which radiate from the opening. A completely closed pore appears as a tiny dense body surrounded by a tight circle of densities at the center of the radiating lines (Fig. 7F). This characteristic structure then disappears, leaving no visible sign of the former pore during interludes.

At the end of a defecation with a single cycle, or after the last cycle of a defecation with multiple cycles, the pore closes completely and disappears. Pore closings between multiple cycles of a defecation are usually complete (13 of 19 or 68%), but sometimes the pore only closes partially (6 of 19 or 32%).

Opening and closing velocities of a given pore are not the same. The fastest speeds of increases in pore diameter during opening were 2.5±0. 52 μm s^−1^ (n = 7 different cycles). The fastest speeds of decreases in pore diameter during closing were 6.4±1.6 μm s^-1^ (n= 6 different cycles). The speed of pore closing is therefore 2-3 times greater than the speed of pore opening. An anal pore is thus not permanently visible by light microscopy in *Mnemiopsis*. Instead, the pore displays a transitory structure that appears at defecation and disappears during interludes.

### Duration of defecations

The duration of a defecation increased with body size. Durations ranged from 2–3 min for larvae and young individuals up to ~2 cm long, and 4–6 min for adults 3–5 cm in body length.

### One functional anal canal and pore

A new and striking feature of successive defecation cycles observed in all animals was that defecations always occurred from the same anal canal/fork in a given animal. No cases of both anal canals/forks defecating, or defecations switching from one anal canal/fork to the opposite one, were observed. This constancy was observed in more than a dozen repeated defecations by individual animals lasting up to seven hours. Main (1928) briefly reported a similar finding in *Mnemiopsis* based on only several defecations.

To test whether only one of the two opposing anal canals/forks is used for defecation in a given animal, I noted which anal canal/fork (right or left) initially defecates in an adult animal pinned in the sagittal plane. The animal was then unpinned, turned over to the other sagittal side, and re-pinned so that the viewer’s right and left anal canal/fork were reversed. Subsequent defecations occurred in the opposite anal canal/fork (i.e., left if previously right and vice versa). Thus, the anal canal/fork which defecates is intrinsic to an animal, independent of which sagittal side is pinned against the Sylgard.

I then tested longer-term constancy of defecation by the same anal canal and fork. To provide an easily visible marker for a defecatory anal canal/fork, a region of the subsagittal comb rows was removed on the side of initial defecation (by right or left anal canal/fork) in 12 animals, which were then unpinned and maintained in running sea water. The wounds healed rapidly and produced easily visible wide gaps between adjacent comb plates on the operated side (Tamm, 2012a). The surgically marked animals were removed daily and pinned in a Sylgard bowl to observe defecation for the next 10 days. In all animals, the anal canal/fork used initially for defecation remained the defecating one. *Mnemiopsis* thus uses the same anal canal/fork and its pore to defecate for at least 10 days.

### Type of diagonal

In aboral views, the defecating and non-defecating major anal lobes lie on a diagonal to the sagittal and tentacular planes. Is this diagonal always the same type, i.e., with a positive slope (bottom left to top right) or with a negative slope (top left to bottom right), or are both types of diagonals found in different animals?

The chondrophore *Velella* possesses a well-known mirror image dimorphism of its diagonal sail, called right-handed or left-handed forms with respect to the downwind sailing direction (Mackie, 1962). Does *Mnemiopsis* possess a similar mirror image dimorphism in the diagonal placement of its major anal lobes?

As described above, the major lobe of one of the two anal canals is larger than the opposite one during interludes and especially at defecations. The type of any diagonal is independent of 180° rotation, but the location of a specific end of a diagonal is reversed by 180° rotation (i.e., for a positive diagonal, top right becomes bottom left). Aboral views of the anal complex in living thick transverse slices from 112 individuals ranging 2–6 cm in body length showed in every case that a greatly swollen defecating lobe with a dilated anal pore expelling waste, or the larger anal lobe of interlude animals, was either at the top right or the bottom left (i.e., near one end of a positive diagonal). Aboral views of eight larvae ranging 2-7 mm long showed six larvae with the defecating pore or the larger interlude anal lobe at the top right or bottom left (i.e., on the positive diagonal as in adults; Figs. 7, 8A). Surprisingly, in two larvae the defecating pore or larger interlude anal lobe was at the top left or bottom right (i.e., on the negative diagonal; Fig. 8B).

**Figure 8.**
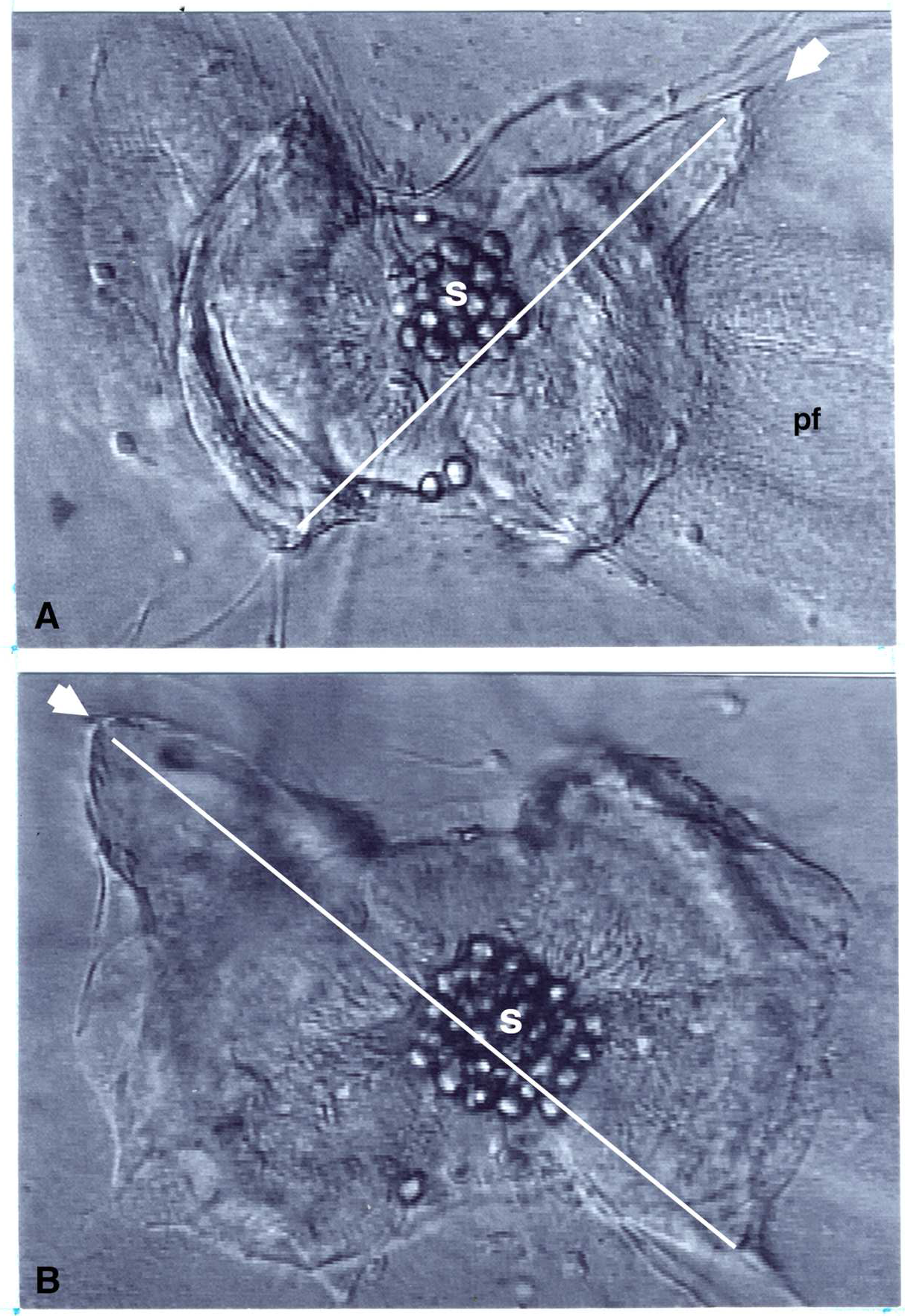
Mirror image dimorphism of the diagonal anal complex in interlude larvae (aboral DIC view, sagittal axis horizontal). No anal pore is visible. A. The 0.18 cm larva of Fig. 7, but during interlude. The larger major anal lobe is at top right (white arrow) on the positive diagonal (white line) intersecting the statocyst (s). pf, polar field on right side. B. A 0.35 cm larva with the larger major lobe at top left (white arrow) on the negative diagonal (white line).

Thus, no mirror image dimorphism of the diagonal anal complex was found in a large sample of adult individuals examined. In a much smaller sample of larvae, the anal pore was on the positive diagonal in most cases but in two cases the pore was on the negative diagonal. Diagonal anal dimorphism thus occurs in larvae but apparently not in adults.

### Defecation rhythm

Defecation occurs rhythmically at regular time intervals in both larvae maintained on microscope slides and in older animals pinned in Sylgard bowls (Table 1, Fig. 9). The time intervals between repeated defecations are generally shorter in smaller animals and longer in larger animals. The frequency of defecation is thus roughly the inverse of body size (Table 1, Fig. 9). Time intervals between defecations in individual animals were remarkably uniform regardless of animal size, conditions of observation (pinned or on slides), or the length of observation. Many of the individuals had been feeding before collection, since remains of ingested prey were commonly seen in the stomodaeum and esophagus.

**Table 1.** Periodicity of defecation in *Mnemiopsis leidyi*, Individuals are listed by increasing size (body length). Successive interval lengths are shown in order from the first to the last defecation observed, along with the mean interval length (± 1 SD). Individuals 2 and 6 had their defecating anal lobe and pore on the negative diagonal (see text). Defecation intervals for individuals 9, 14 and 23 are shown in Fig. 9.

| Individual | Size (cm) | Interval lengths (min) | Mean interval length (min) |
| --- | --- | --- | --- |
| 1 | 0.18 | 18, 17, 21, 24, 25 | 21 ± 3.5 |
| 2 | 0.20 | 20, 16, 14 | 17 ± 3.1 |
| 3 | 0.20 | 12, 15 | 14 ± 2.1 |
| 4 | 0.25 | 12, 15, 13, 16 | 14 ± 1.8 |
| 5 | 0.30 | 25, 25, 26, 23, 21, 23, 20, 22 | 23 ± 2.1 |
| 6 | 0.35 | 16, 15, 18, 18, 19, 22 | 18 ± 2.4 |
| 7 | 0.40 | 25, 25, 31, 27 | 27 ± 2.8 |
| 8 | 0.55 | 29, 19, 19 | 22 ± 5.8 |
| 9 | 0.60 | 10, 10, 11, 10, 12, 11, 8, 10, 11, 11 | 10 ± 1.1 |
| 10 | 0.70 | 19, 11, 13, 14, 24, 18, 21 | 17 ± 4.7 |
| 11 | 1.2 | 30, 30, 27, 28, 27, 29 | 29 ± 1.4 |
| 12 | 1.2 | 17, 17, 17, 18 | 17 ± 0.5 |
| 13 | 1.3 | 30, 32, 26, 28, 24, 24, 30, 29 | 28 ± 2.9 |
| 14 | 1.3 | 21, 21, 20, 28, 27, 23, 23, 19, 20, 21, 21 | 22 ± 2.9 |
| 15 | 1.3 | 27, 25, 24, 36 | 28 ± 5.5 |
| 16 | 1.5 | 35, 27, 28, 38, 40 | 34 ± 5.9 |
| 17 | 1.7 | 37, 28, 26, 34, 27, 26, 31, 32, 30, 37, 29, 26, 34 | 31 ± 4.0 |
| 18 | 2.0 | 35, 25, 25, 23 | 27 ± 5.4 |
| 19 | 2.2 | 37, 45 | 41 ± 5.7 |
| 20 | 2.2 | 34, 43, 47 | 41 ± 6.7 |
| 21 | 3.0 | 30, 40, 43, 42, 35 | 38 ± 5.4 |
| 22 | 3.0 | 35, 40, 37 | 37 ± 2.5 |
| 23 | 4.0 | 54, 56, 55, 58 | 56 ± 1.7 |
| 24 | 4.2 | 57, 46, 43, 49, 53 | 50 ± 5.5 |
| 25 | 4.5 | 51, 57, 42, 48, 50, 47 | 49 ± 5.0 |
| 26 | 5.0 | 45, 45, 45, 45 | 45 ± 0 |
| 27 | 5.0 | 56, 55, 56, 61 | 57 ± 2.7 |
| 28 | 5.0 | 60, 52, 58, 63, 57 | 58 ± 4.1 |

**Figure 9.**
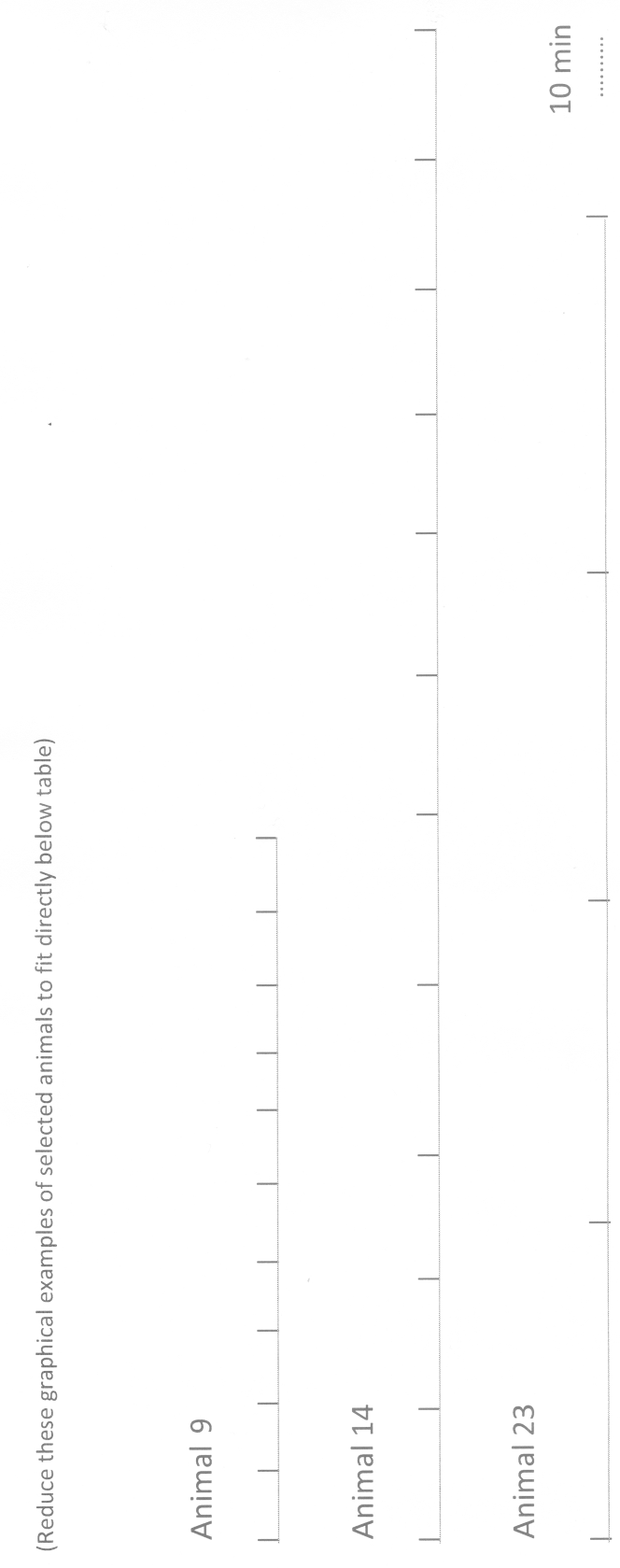
Graphical examples of intervals between defecations in selected animals of different sizes: animal 9 (0.6 cm), animal 14 (1.3 cm) and animal 23 (4.0 cm) are from Table 1. Tick marks indicate onset of defecations. Note that the interval between successive defecations increases with body size.

## Discussion

Ever since the classic studies by Agassiz (1850) and Chun (1880), it has been well documented that ctenophores do not possess a bottle-gut, but have a through-gut with mouth and separate anus (Tamm, 2016), as confirmed again by Presnell et al, 2016. However, many important aspects of the gastrovascular system in ctenophores have escaped notice or were incorrectly reported. I show here that the process of defecation in *Mnemiopsis* occurs with an ultradian rhythm using only a single anal pore as the anus. Contrary to the literature, the anal pore of *Mnemiopsis* is not visible by light microscopy during interludes, but appears at defecation and disappears afterward.

A basic premise of this study is that changes in the width and flow patterns of endodermal canals play a major role in food distribution and waste elimination. Stereo and DIC microscopy demonstrate that changes in canal diameter are not due to attached mesogleal muscles, as reported by Presnell et al (2016). Nor have muscle cells or myoendodermal cells been found within the canal walls themselves (Hernandez-Nicaise, 1991; pers. comm., 2016)

Widening and narrowing of the canals are more likely due to fluctuations in hydrostatic pressure within the canal lumens by fluid flow driven by cilia lining their walls. Franc (1972) found in *Beroe ovata* and *Leucothea multicornis* that an increase in internal pressure causes progressive expansion of endodermal canals, leading to swelling of anal canals and opening of anal pores to expel waste products. The venting of fluid was found to restore normal pressure and diameter of canals, leading to closing of anal pores. Injecting sea water into meridional canals to artificially increase internal pressure in the gastrovascular system produced the same events (Franc, 1972). These experiments indicate that swelling and narrowing of endodermal canals as well as opening and closing of anal pores are due to changes in hydrostatic pressure within the gastrovascular system. Changes in shape of the stomach and corrugation of the esophageal walls in *Mnemiopsis*, however, are probably due to intracellular contractile elements within these organs.

A major unsolved problem is how the ciliary motor responses underlying the patterns of observed fluid and particle flow in the gastrovascular system are initiated and controlled during interludes and defecations. Recent immunofluorescence staining of the nervous system of *Pleurobrachia pileus* has not revealed an endodermal nerve net (Jager et al., 2011). Conduction pathways mediating coordination and control of gastrovascular responses in *Mnemiopsis* remain a fruitful subject for future investigations.

### The dynamic anal pore

Video imaging of defecation in *Mnemiopsis* revealed that a new pore suddenly appears and opens between the closely apposed endodermal and ectodermal layers at the tip of the protruding major anal lobe, thereby establishing continuity with the external sea water and allowing exit of fecal matter from anal canals. After defecation the pore closes and is not visible during interludes, contrary to their ubiquitous depiction in non-defecating ctenophores in the literature.

This “now-you-see-it-now-you-don’t” performance by the anal pore in *Mnemiopsis* is puzzling. Does the pore really disappear completely, i.e., by disassembly and/or breakdown of its components? Or is the closed pore in its final constricted state simply too small to see by the DIC video imaging system used here? The former possibility represents a novel ephemeral pore that rhythmically is lost and regenerates at every defecation. If so, defecation periodicity and duration indicates that *Mnemiopsis* would spend ~90% of its life without an anal structure or through-gut, and thus have an intermittent anus and intermittent through-gut. Such temporality of the anus might provide clues for understanding evolution of a permanent anus and through-gut in animals. The latter possibility of optical limitations hiding a closed pore argues for a permanent anal pore as found in all other animals. Further investigations using more incisive approaches are required to settle this matter (see below).

The mechanism of anal pore function is also puzzling. By DIC microscopy of working anal pores, the enlarging rim of an opening pore resembles an expanding ring of endodermal-ectodermal fusion, and the decreasing rim of a closing pore looks like a diminishing ring of transepithelial fusion. Such a mechanism, if confirmed by further investigations, would be the first known example of different germ layers locally and reversibly melding and separating to form transitory spatial continuities for a physiological function. By contrast, the anus of other protostomes is a permanent organ formed by stable fusion of endodermal and ectodermal epithelia at the posterior end of the embryo.

Fluorescence staining of F-actin (Presnell et al, 2016) and myosin II heavy chains (Dayraud et al, 2012) in the presumed location of anal pore(s) in fixed individuals of *Mnemiopsis* and *Pleurobrachia* were taken as evidence that anal pores function in a more conventional manner as permanent ring-like contractile sphincters (Presnell et al, 2016). Whether DIC microscopy of living anal pores would necessarily discern the formation of an actin-based sphincter is not clear. Polarized light microscopy that detects weak birefringence might be more suitable, and perhaps a fluorogenic probe for vital staining of F-actin, such as SiR-actin, would be useful. But ultrastructural methods using high pressure frozen tissues are probably the most conclusive way to clarify these questions about anal pore existence and operation. Ideally, this approach would capture for TEM identified stages of pore opening, closing and disappearance to image endodermal and ectodermal epithelial cells during pore function.

Franc’s (1972) experiments and the present findings indicate that the appearance of the anal pore is initiated by elevated hydrostatic pressure that causes anal lobe protrusion and close apposition of endodermal and ectodermal tissue at the tip, leading to local epithelial fusion to form an enlarging pore. Pore closing and disappearance would then result from a drop in internal pressure after fluid expulsion, leading to separation of the endodermal and ectodermal epithelial layers and withdrawal of the lobe from the aboral surface. Ciliary fluid flow may thus regulate hydrostatic pressure in the gastrovascular system and effect the dynamics of the anal pore.

### Only one functional anal canal and pore in *Mnemiopsis*

This study shows that *Mnemiopis* uses only one of their two bi-lobed anal canals (forks) to defecate. Except for a brief mention to this effect by Main (1928), all previous studies of defecation in ctenophores, from Agassiz (1850) and Chun (1880) to Presnell et al (2016), report that both diagonally opposite major lobes of the two anal canals have anal pores which defecate, i.e., two functional anal canals and pores (Fig. 10B).

**Figure 10.**
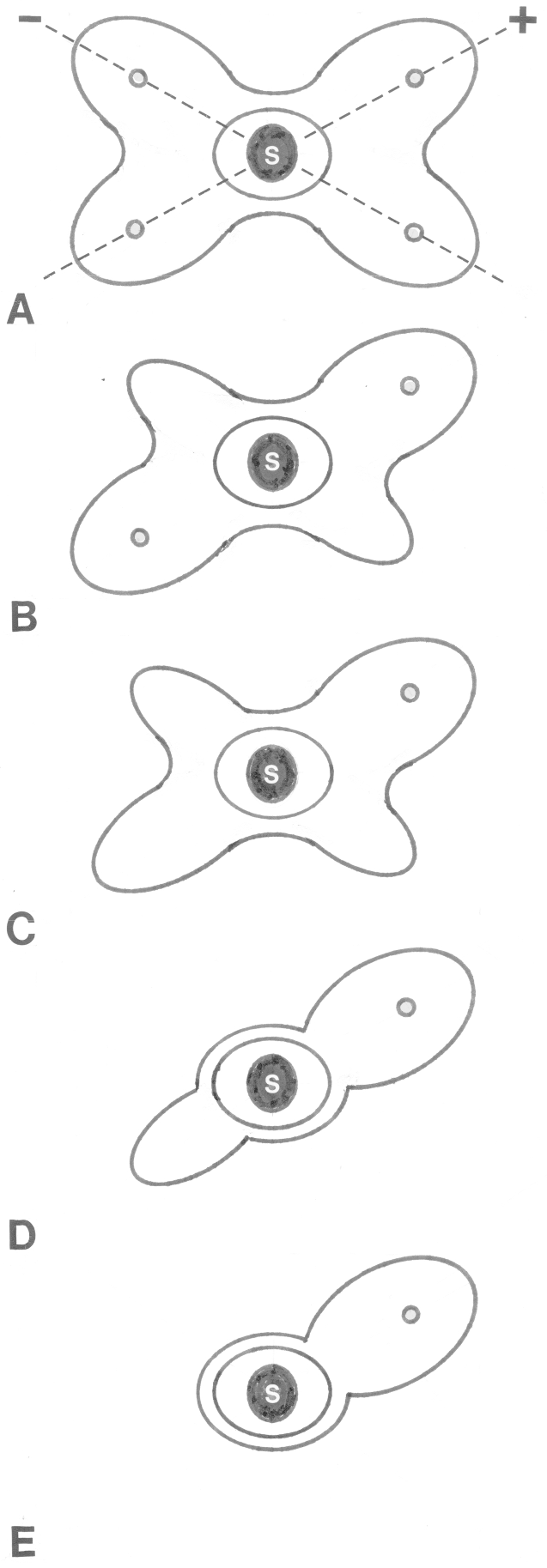
Hypothetical evolutionary sequence of patterns of interlude anal architecture. Positive (+) and negative (−) diagonals intersect the statocyst (s). For convenience, pore locations during defecation are indicated here. A. Possible past or future pattern of four equivalent defecatory major lobes with four pores. B. Widely accepted pattern (Hyman, 1940) of two equivalent defecatory major lobes with two pores, and two blind minor lobes. C. Pattern found here: one larger defecatory major lobe with the single anal pore, a smaller non-defecatory major lobe without a pore, and two blind lobes. D. Possible past or future stage of one defecatory major lobe and one opposite blind lobe. E. Possible first or last stage in this series with one defecatory major lobe and pore only.

I believe this discrepancy may be due to several factors. Firstly, the diagonally opposite major lobes of the two anal canals look superficially alike at defecation when both major lobes swell and become filled with waste particles. In randomly oriented *Mnemiopsis*, either anal canal (right or left fork in sagittal views; top right or bottom left lobe in aboral views) is as likely as the other one to be used for defecation in an individual animal, as found here. It is not surprising that Agassiz (1850) and Chun (1880) believed that both diagonally opposed major anal lobes had pores which defecated. However, these authors never described two anal pores defecating at the same time. Nor did Agassiz or Chun describe individual animals switching between alternate pores to defecate. Without following each anal canal/fork during repeated defecations of the same animal as done here, previous workers may have mistakenly presumed similar anal functions for similar anal structures. Agassiz’s and Chun’s descriptions were adopted by Hyman (1940) in her authoritative textbook, which has served as the basis for anal anatomy and function in ctenophores (Fig. 10B).

A second reason for the discrepancy between my findings and previous studies is that the pattern of defecation found in *Mnemiopsis* may not occur in all ctenophores. In fact, preliminary observations on *Pleurobrachia pileus* from Woods Hole, MA reveal that both major anal lobes have pores which defecate (Tamm, in preparation). The pattern of defecation in *Mnemiopsis* may thus not be common to all ctenophores. Comparative investigations of defecation in various types of ctenophores, following the methods used here, are needed to resolve this matter.

Nevertheless, the structural and functional difference found here between the diagonally opposed major anal lobes is an intriguing exception to the 180° rotational symmetry of ctenophores (Dunn et al., 2015). This discovery provides a unique body marker for sidedness in the symmetry plan of *Mnemiopsis* and perhaps other ctenophores.

### Diagonal types

The present finding that the major anal lobes lie on the positive diagonal of the anal complex in all adult *Mnemiopsis* and most larvae confirms descriptions of other ctenophores by Chun (1880), Mayer (1912) and Hyman (1940). It was therefore surprising that in two larvae the diagonal of the major anal lobes was the negative type. This mirror image anal dimorphism in several larvae may be related to developmental flexibility in symmetry of the anal complex that for some reason becomes less common in adults.

### Defecation rhythm

The regular periodicity of defecation found in *Mnemiopsis* (Table 1; Fig. 9) is the first documented example of an ultradian rhythm in the phylum Ctenophora. In contrast, diverse types and functions of ultradian rhythmic behaviors are common and well-studied in Cnidaria (Passano, 1963; Bullock and Horridge, 1965; Anderson, 1985; Mackie, 1980;
Satterlie, 1985).

The relative importance of endogenous and exogenous factors in the onset, production and regulation of biological rhythms is a well-studied topic. The rhythmic defecations in *Mnemiopsis* appear to originate within the animal, since they continue under uniform laboratory conditions without feeding or known external stimuli. However, many of the animals had been eating before being collected, so feeding may be an exogenous factor promoting onset of rhythmic defecation. It will be important to test effects of starvation on the initiation and frequency of defecation. The finding that the periodicity of defecation depends roughly on animal size also suggests an endogenous origin of rhythmicity.

### Anal considerations

An interesting question is why the ctenophore gut does not run straight through the body to the anus, but is branched at the anal end. The aboral pole is instead occupied by the statocyst in the apical organ. I believe this may be an evolutionary outcome of competition for the aboral site between two important functional systems: geotaxis vs. defecation by a through-gut. A single statocyst that regulates the activity of eight locomotory comb rows responsible for geotactic behavior would not work effectively if it were located off the aboral-oral body axis (Tamm, 1982, 2014a, 2014b, 2015). An anus, on the other hand, should not be subject to off-center functional constraints. The statocyst, according to this argument, had evolutionary priority over the anus for first place at the aboral (south) pole. During embryonic development, the apical organ arises before the anal complex forms (Mayer, 1912), thereby setting the stage for its later dominance at the aboral end.

It is more difficult to understand the pattern of anal canal branching and anal pore number and location around the apical organ (Fig. 10). In contrast to the widely accepted view of two equal major anal lobes and two pores (Fig. 10B), I found that *Mnemiopsis* has two unequal major lobes, one of which is larger than the other and hosts the single anal pore at defecation (Fig 10C). What is the present-day functional reason for this asymmetry? Is this pattern a secondary reduction or loss from an earlier stage with four equivalent anal canals with four pores on both diagonals (Fig. 10A)? Or is the existing pattern an intermediate step in the reverse direction from a simpler stage (Fig. 10E) to a full-blown anal foursome (Fig. 10A)? Unfortunately, fossil records of ctenophores are disputable (Steve Haddock, MBARI, pers. comm., 2014) and directions of natural evolution are dialectic and unpredictable.

## Acknowledgments

This research was supported by a gift from the Fund for Science at MBL. I thank Mark Terasaki, Laurinda Jaffe, John Pearse, Vicki Pearse, Gary Freeman and Steve Haddock for helpful discussion. Comments by the reviewers enhanced the paper. Molly McQuillan in the Bell Center at MBL kindly scanned the figures.

## References

Agassiz L 1850. Contributions to the natural history of the Acalephae of North America. Amer. Acad. Arts Sci., Part 2: 313–374.

Anderson PAV 1985. The comparative electrobiology of gelatinous zooplankton. Bull. Mar. Sci. 37: 460–477.

Barnes RD 1980. Invertebrate Zoology. Fourth Edition. Philadelphia: Saunders/Holt, Reinhart and Winston.

Brusca RC & Brusca GJ 1990. Invertebrates. Oxford, UK: Sinauer Associates.

Bullock TH & Horridge GA 1965. Structure and Function in the Nervous Systems of Invertebrates. Vol. I. San Francisco: W. H. Freeman.

Bumann D & Puls G 1997. The ctenophore Mnemiopsis leidyi has a flow-through system for digestion with three consecutive phases of extracellular digestion. Physiol. Zool. 70: 1–6.

Chun C 1880. Die Ctenophoren des Golfes von Neapel und der angrenzenden Meeres-Abschnitte. Flora und Fauna des Golfes von Neapel, Vol. 1. Leipzig: Engelmann.

Colin SP, Costello JH, Hansson LJ, Titelman J & Dabiri JO 2010. Stealth predation and the predatory success of the invasive ctenophore Mnemiopsis leidyi. Proc. Natl. Acad. Sci. USA 107: 17223–17227.

Dayraud C, Alie A, Jager M, Chang P, Le Guyader H, Manuel M & Queinnec E 2012. Independent specialisation of myosin II paralogues in muscle vs. non-muscle functions during early animal evolution: a ctenophore perspective. BMC Evol. Biol. 12: 107–128.

Dunn CW, Leys SP & Haddock SHD 2015. The hidden biology of sponges and ctenophores. Trends Ecol. Evol. 30: 282–291.

Dunn CW & Ryan JF 2015. The evolution of animal genomes. Curr. Opin. Genetics and Development. 35: 25–32.

Franc JM 1972. Activités des rosettes ciliées et leur supports ultrastructuraux chez les Cténaires. Z. Zellforsch. 130: 527–544.

Gemmill JF 1918. Ciliary action in the internal cavities of the ctenophore Pleurobrachia pileus Fabr. Proc. Zool. Soc. Lond. 1918: 263–265.

Haddock SHD 2007. Comparative feeding behavior of planktonic ctenophores. Integr. Comp. Biol. 47: 847–853.

Harbison GR, Madin LP & Swanberg NR 1978. On the natural history and distribution of oceanic ctenophores. Deep-Sea Res. 25: 233–256.

Harbison GR & Madin LP 1982. Ctenophora. In Synopsis and Classification of Living Organisms. (ed. S. P. Parker), pp. 707–717. New York: McGraw-Hill.

Hernandez-Nicaise M-L 1991. Ctenophora. In Microscopic Anatomy of Invertebrates. Vol. 2. Placozoa, Porifera, Cnidaria, and Ctenophora. (ed. F. W. Harrison and J. A. Westfall), pp. 359–418. New York: Wiley-Liss.

Hyman LH 1940. The Invertebrates: Protozoa through Ctenophora. Vol. 1. New York: McGraw-Hill.

Jager M, Chiori R, Alie A, Dayraud C, Queinnec E & Manuel M 2011. New insights on ctenophore neural anatomy: Immunofluoresence study in Pleurobrachia pileus (Muller, 1776). J. Exp. Zool. (Mol. Dev. Evol.) 316: 171–187.

Mackie GO 1962. Factors affecting the distribution of Velella (Chondrophora). Int. Revue des Hydrobiol. 47: 26–32.

Mackie GO 1980. Jellyfish neurobiology since Romanes. Trends Neurosci. 3: 13–16.

Main RJ 1928. Observations of the feeding mechanism of a ctenophore, Mnemiopsis leidyi. Biol. Bull. 55: 69–78.

Mayer AG 1912. Ctenophores of the Atlantic Coast of North America. Washington, D. C.: Carnegie Institution of Washington.

Moss AG 1991. The physiology of feeding in the ctenophore Pleurobrachia pileus. Hydrobiologia 216: 19–25.

Passano LM 1963. Primitive nervous systems. Proc, Natl. Acad. Sci. USA 50: 306–313.

Pearse V, Pearse J, Buchsbaum M & Buchsbaum R 1987. Living Invertebrates. Palo Alto, CA: Blackwell Scientific and Pacific Grove, CA: Boxwood Press.

Presnell JS, Vandepas LE, Warren KJ, Swalla BJ, Amemiya CT & Browne WE 2016. The presence of a functionally tripartite through-gut in ctenophores has implications for metazoan character trait evolution. Curr. Biol. 26: 1–7.

Satterlie RA 1985. Central generation of swimming activity in the hydrozoan jellyfish Aequorea aequorea. J. Neurobiol. 16: 41–55.

Sullivan LJ 2010. Gut evacuation of larval Mnemiopsis leidyi A. Agassiz (Ctenophora, Lobata). J. Plankton Res. 32: 69–74.

Swanberg N 1974. The feeding behaviour of Beroë ovata. Mar. Biol. 24: 69–76.

Tamm SL 1982. Ctenophora. In Electrical Conduction and Behaviour in ‘Simple’ Invertebrates. (ed. G. A. B. Shelton), pp. 266–358. Oxford, UK: Oxford University Press.

Tamm SL 2012a. Regeneration of ciliary comb plates in the ctenophore Mnemiopsis leidyi. I. Morphology. J. Morphol. 273: 109–120.

Tamm SL 2012b. Patterns of comb row development in young and adult stages of the ctenophores Mnemiopsis leidyi and Pleurobrachia pileus. J. Morphol. 273: 1050–1063.

Tamm SL 2014a. Cilia and the life of ctenophores. Invertebr. Biol. 133: 1–46.

Tamm SL 2014b. Formation of the statolith in the ctenophore Mnemiopsis leidyi. Biol. Bull. 227: 7–18.

Tamm SL 2015. Functional consequences of the asymmetric architecture of the ctenophore statocyst. Biol. Bull. 229: 173–184.

Tamm SL 2016. No surprise that comb jellies poop. Science 352: 1182.

Tamm SL & Moss AG 1985. Unilateral ciliary reversal and motor responses during prey capture by the ctenophore Pleurobrachia. J. Exp. Biol. 114: 443–461.

Tamm SL & Tamm, S 1987. Massive actin bundle couples macrocilia to muscles in the ctenophore Beroë. Cell Motil. Cytoskeleton 7: 116–128.

Tamm SL & Tamm, S 1991. Reversible epithelial adhesion closes the mouth of Beroë, a carnivorous marine jelly. Biol. Bull. 181: 463–473.

Waggett R & Costello JH 1999. Capture mechanisms used by the lobate ctenophore Mnemiopsis leidyi preying on the copepod Acartia tonsa. J. Plankton Res. 21: 2037–2052.

